# Perception of effort during an isometric contraction is influenced by prior muscle lengthening or shortening

**DOI:** 10.1101/2021.03.29.437599

**Authors:** Benjamin Kozlowski, Benjamin Pageaux, Emma F. Hubbard, Benjamin St. Peters, Philip J. Millar, Geoffrey A. Power

**Author notes:** Correspondence:* Geoffrey A Power PhD., **N**euromechanical **P**erformance **R**esearch **L**aboratory, Department of Human Health and Nutritional Sciences, College of Biological Sciences, University of Guelph, Ontario, Canada.

## Abstract

**Purpose:** Following a shortening or lengthening muscle contraction, the torque produced in the isometric steady state is distinctly lower (residual torque depression; rTD) or higher (residual torque enhancement; rTE), respectively, compared to a purely isometric contraction at the same final muscle length and level of activation. This is referred to as the history dependence of force. When matching a given torque level, there is greater muscle activation (electromyography; EMG) following shortening and less activation following lengthening. Owing to these differences in neuromuscular activation, it is unclear whether perception of effort is altered by the history dependence of force.

**Methods:** Experiment 1 tested whether perception of effort differed between the rTD and rTE state when torque was matched. Experiment 2 tested whether perception of effort differed between the rTD and rTE state when EMG was matched. Finally, experiment 3 tested whether EMG differed between the rTD and rTE state when perception of effort was matched.

**Results:** When torque was matched, both EMG and perception of effort were higher in the rTD compared to rTE state. When EMG was matched, torque was lower in the rTD compared to rTE state while perception of effort did not differ between the two states. When perception of effort was matched, torque was lower in the rTD compared to rTE state and EMG did not differ between the two states.

**Conclusion:** The combined results from these experiments indicate that the history dependence of force alters one’s perception of effort, dependent on the level of motor command.

## Introduction

Following active shortening and lengthening contractions, steady state isometric torque is reduced or elevated, respectively, compared to an isometric contraction at the same final muscle length and level of activation without prior shortening or lengthening (Abbott and Aubert, 1952; Altenburg et al., 2008; Herzog, 2004, 2000; Lee et al., 2001). These two phenomena are referred to as residual torque depression (rTD) and residual torque enhancement (rTE), respectively (Herzog, 2004), and have been observed in humans at the single muscle fibre level (Pinnell et al., 2019), and across multiple muscle groups and contraction intensities during voluntary and electrically stimulated contractions (Chapman et al., 2018; Chen et al., 2019; Seiberl et al., 2015). However, it is unclear whether this history-dependent phenomena affects perception of effort.

Owing to the history dependence of force, there is greater neuromuscular activation as assessed with electromyography (EMG) in the rTD state following active shortening (activation increase) and lower activation in the rTE state following active lengthening (activation decrease) relative to a purely isometric contraction performed at the same absolute torque (Herzog, 2004; Jones et al., 2016; Kosterina et al., 2013; Mazara et al., 2018; Seiberl et al., 2015). As neuromuscular activation assessed by EMG is traditionally used as a proxy of motor command (Carrier et al., 2011; Thoroughman and Shadmehr, 1999), differences in EMG caused by the history dependence of force may indicate alterations in motor command during the rTD and rTE states. Seed et al. (2018) recently confirmed these observations, but also demonstrated that during an isometric contraction performed at the same absolute torque in the rTD and rTE state, heart rate and blood pressure responses were higher in the rTD state compared to the rTE state. As heart rate and blood pressure responses are two parameters known to be influenced, at least in part, by motor command (Engel and Talan, 1991; Gallagher et al., 2001; Gandevia and Hobbs, 1990; Matsukawa et al., 2013, 2007; Murata et al., 2004; Victor et al., 1989), these results provide direct evidence that the rTD and rTE states can influence motor command. However, it is less clear how rTD and rTE influence perception of effort.

Perception of effort, defined as the conscious sensation of how hard, heavy, and strenuous a physical task is (Marcora, 2009; for review see: Pageaux, 2016), has been proposed to be related to the magnitude of the motor command sent to the working muscles (de Morree et al., 2014, 2012; Pageaux & Gaveau, 2016; Siemionow et al., 2000). Considering that the rTD state increases objective markers of the motor command (heart rate, blood pressure, and EMG) during an isometric contraction and rTE decreases these markers, an investigation of the relationship between the magnitude of the motor command sent to the working muscles and the perception of effort, as well as how the history dependence of force alters one’s perception of effort is warranted.

The purpose of the current study was to assess whether the history dependence of force can alter one’s perception of effort. In experiment 1, participants performed a torque matching protocol to control for motor output. EMG and perception of effort were measured while the participants performed an isometric contraction at the same absolute torque level preceded by either a shortening (rTD state) or lengthening (rTE state) contraction. We expected that for the same absolute torque, EMG and perception of effort would be higher in the rTD compared to the rTE state. In experiment 2, participants performed an activation (EMG) matching protocol to control for the magnitude of the motor command by using EMG as an objective marker of the motor command sent to the working muscle (Carrier et al., 2011; Thoroughman and Shadmehr, 1999). Torque and perception of effort were measured while the participants performed an isometric contraction at the same absolute torque preceded by either a shortening (rTD state) or lengthening (rTE state) contraction. We expected that for the same level of EMG, the torque produced would be lower in the rTD compared to the rTE state, with no difference in perception of effort. In experiment 3, participants performed a perceived effort matching protocol to control for the magnitude of the motor command by using perception of effort as a subjective marker of the motor command sent to the working muscle (de Morree et al., 2012; Gandevia et al., 1981; Gandevia and McCloskey, 1977; Kernell, 1995; Kim et al., 2004). Torque and EMG were measured while the participants performed an isometric contraction at the same intensity of perceived effort preceded by either a shortening (rTD sate) or lengthening (rTE state) contraction. We expected that for the same level of effort, the torque produced would be lower in the rTD compared to the rTE state, with no difference in EMG. These expected results would provide additional support to the hypothesis that the history dependence of force (rTD vs rTE state) influences the magnitude of the motor command required to perform an isometric contraction and effectively alters one’s perception of effort.

## Methods

### Participant Information

Twenty-two male participants (age: 24.2 ± 2.1 yrs., height: 176.4 ± 7.5 cm, weight: 78.2 ± 13.1 kg) participated in experiment 1, and fifteen male participants (age: 22.3 ± 2.5 yrs., height: 178.2 ± 6.0 cm, weight: 77.0 ± 9.2 kg) participated in experiment 2 and experiment 3. All participants were recreationally active and free of any known cardiovascular or neuromuscular diseases. Participants were excluded from the study if they self-reported any physical or mental fatigue or were taking any medication that would alter heart rate and blood pressure. All participants gave written informed consent and all procedures were approved by the Research Ethics Board of the University of Guelph.

### Participant Setup for Experimental Protocols

A HUMAC NORM dynamometer (CSMi Medial Solutions, Stoughton, MA) was used for all torque, angular velocity, and joint position recordings. All testing and data collection were performed on the right lower limb. Each participant was seated on the dynamometer with their right hip at 110° of flexion and right knee at 140° of flexion (180°; straight). The right knee was immobilized with an inelastic strap, positioned superior to the right patellar tendon on the distal-anterior thigh. A cushioned pad was located underneath the right thigh to properly position the knee and ankle joints. Movement of the torso was restricted with a four-point seatbelt harness. The right foot was fixed to a dorsi-/plantar flexor dynamometer pedal adaptor with one inelastic strap placed over the ankle at the level of the lateral and medial malleoli, and another at the distal portion of the metatarsals. The minimum and maximum ankle plantar flexion angles were set to 10° dorsiflexion and 50° plantar flexion, with the isometric reference angle set at 20° plantar flexion (all relative to a neutral ankle angle at 90°; 0° reference). This setup allowed for a 30° ankle rotation during muscle shortening and lengthening relative to the 20° isometric plantar flexion angle, allowing all experimental conditions to be assessed at the same final joint angle throughout the testing protocols.

### Electromyographic Placement for Recording

Neuromuscular tests are represented in Figure 1. Prior to electromyography (EMG) electrode placement, skin locations were thoroughly shaven, and cleaned with alcohol swabs. Silver-silver chloride (Ag/AgCl) electrodes (1.5 x 1 cm: Kendall, Mansfield, MA) were placed using a large distance (8 −10 cm) bipolar configuration and used for all EMG recordings. One electrode was placed on the muscle belly of the soleus at the midline of the lower leg, located by palpating the soleus muscle belly approximately 2 cm inferior to the medio-inferior border of the gastrocnemius, while the participant plantarflexed. A reference electrode was placed distally. To record antagonist muscle activation, one electrode was positioned over the muscle belly of tibialis anterior, located by palpating 7 cm distal and 2 cm lateral to the tibial tuberosity, while the participant dorsiflexed. A reference electrode was placed distally. One ground electrode was placed over the patella of the right leg.

**Fig. 1.**
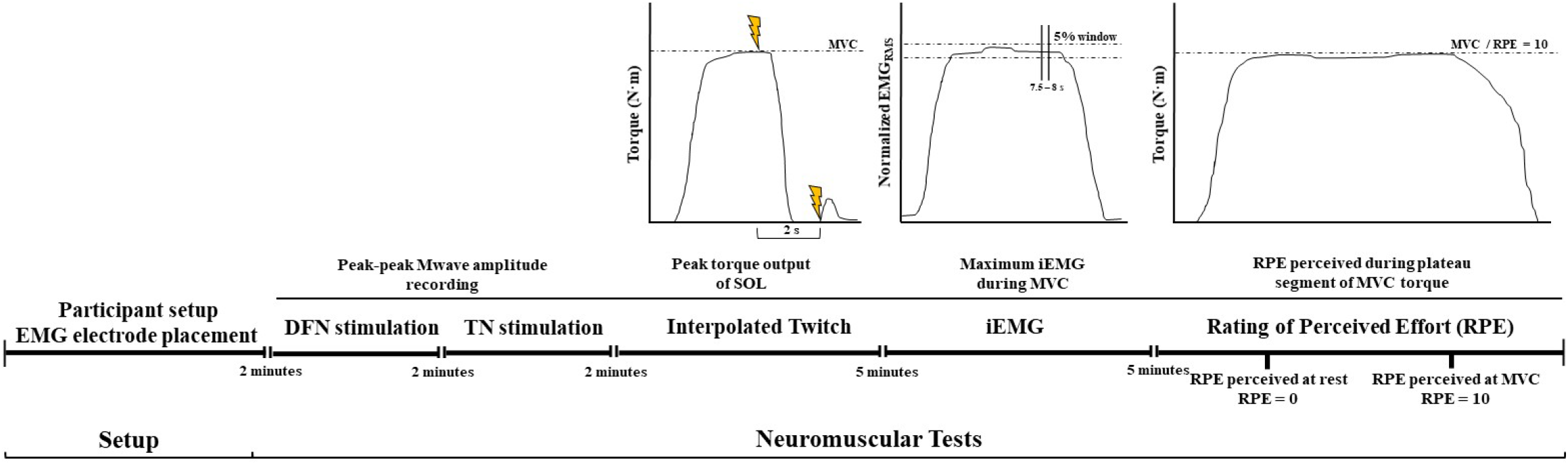
Pre-experimental protocol overview. Setup consisted of participant positioning on the dynamometer seat and surface EMG electrode placement. Neuromuscular testing followed setup. First, deep fibular nerve (DFN) stimulation followed by tibial nerve (TN) stimulation were conducted to evoke appropriate M-waves. Following peripheral nerve stimulation, maximal torque (MVC) output was deduced using the interpolated twitch-technique, followed by maximum iEMG magnitude. The final neuromuscular test conducted was used for the anchoring of the rating of perceived effort (RPE) scale. Time intervals represent rest intervals given for participants between protocols.

### Neuromuscular Responses from Deep Fibular and Tibial Nerve Stimulation

Maximal compound muscle action potentials (M-wave) were obtained to normalize participant EMG root mean squared (EMG_RMS_). M-waves were recorded from the soleus and tibialis anterior muscles by stimulating the tibial and deep fibular nerve, respectively, with a standard clinical bar electrode (Empi, St Paul, Minnesota, USA) coated in conductive gel. Participants were at rest during all peripheral nerve stimulation. Peripheral nerve stimulation was performed with a single pulse from a constant current high voltage stimulator (model DS7AH, Digitimer, Welwyn Garden City, Hertfordshire, UK). Voltage was set to 400 V and pulse width to 200 μs. Current increased, starting from 20 mA, until a peak-to-peak M-wave amplitude plateau (M_max_) was reached. The current was adjusted to a supramaximal level, equivalent to 120% of that required to generate the M_max_, ensuring the stimulation of all motor units (Neyroud et al., 2014).

### Determining Maximal Voluntary Activation Level

To determine participants maximal voluntary activation (VA) from the plantar flexor muscles, the interpolated twitch technique was used during a 5 s maximal voluntary contraction (MVC) (Behrens et al., 2017; Cheng et al., 2013). The ankle was set at 20° plantar flexion. A twitch was evoked at the torque plateau during the 5 s MVC (interpolated twitch) and a second one at rest, 1 s after the end of the MVC (resting twitch). VA was assessed as: *(1 – Interpolated Twitch Torque Amplitude / Resting Twitch Torque Amplitude) × 100*. Participants were verbally encouraged during the MVC and torque traces were visible to the participant on a computer monitor. All participants could reach a minimum of 95% VA prior to and following all experiments (Table 1).

**Table 1.** Neuromuscular values of participants in experiment 1 and experiments 2 and 3 obtained during the neuromuscular tests performed prior to experimental protocols

| Neuromuscular Assessment | Experiment 1<br>(n = 22) | Experiments 2 and 3<br>(n = 15) | Independent t-test<br>Result |
| --- | --- | --- | --- |
| Pre- experimental MVC Torque<br>(N·m) | 59.7 ± 19.2 | 61.0 ± 13.9 | p = 0.818 |
| Post-experimental MVC Torque<br>(N·m) | 57.2 ± 11.8 | 59.4 ± 14.4 | p = 0.677 |
| Pre-experimental MVC<br>EMG<sub>RMS</sub><br>(mV) | 0.81 ± 0.61 | 0.80 ± 0.55 | p = 0.863 |
| Post-experimental MVC<br>EMG<sub>RMS</sub><br>(mV) | 0.80 ± 0.43 | 0.78 ± 0.31 | p = 0.979 |
| Pre-experimental VA<br>(%) | 98.6 ± 0.7 | 98.9 ± 0.5 | p = 0.224 |
| Post-experimental VA<br>(%) | 98.8 ± 0.5 | 98.4 ± 1.6 | p = 0.258 |
| SOL M<sub>max</sub><br>(mV) | 25.9 ± 7.0 | 26.9 ± 6.7 | p = 0.611 |
| SOL M<sub>max</sub> EMG<sub>RMS</sub><br>(mV) | 5.87 ± 2.1 | 6.23 ± 1.8 | p = 0.592 |
| TA M<sub>max</sub><br>(mV) | 17.7 ± 7.5 | 16.7 ± 3.6 | p = 0.549 |
| TA M<sub>max</sub> EMG<sub>RMS</sub><br>(mV) | 4.07 ± 1.00 | 4.54 ± 0.81 | p = 0.139 |

Values are represented as means ± SD. Participants in experiment 2 and 3 were the same. There was no difference ( $p > 0.05$ ) in all force-production and peripheral parameter means between the participants completing experiment 1, and experiments 2 and 3.
SOL: Soleus ; TA: Tibialis anterior

### Determining Submaximal Torque and Activation Levels

In each experiment, submaximal targets of torque and EMG were assessed for each participant prior to the experimental protocols and visual target lines to match were provided on a computer monitor. Participant EMG was assessed under two matching submaximal conditions using integrated EMG (iEMG), while recording the torque produced. To determine the submaximal iEMG target, participants were instructed to perform a 10 s maximal isometric plantar flexion contraction at an ankle angle of 20° plantar flexion. The average iEMG collected between 7.5 −8 s was then used to determine the 20% and 80% iEMG targets. A ± 5% window was calculated around the targets, and for all subsequent activation-matched contractions, participants were instructed to maintain their iEMG within guidelines marking this target window. Torque feedback was calculated based on the signal averaged for the same time and window as described above for iEMG.

### Anchoring the Rating of Perceived Effort

Following the assessment of maximal iEMG, participants were briefed on the CR10 Borg Scale (Borg, 1982; Grant et al., 1999; Williams, 2017) by explaining its use to gauge the amount of effort (perception of effort) exerted during a contraction from a rating of perceived effort (RPE) scale of 0 – 10 (0 being no effort and 10 being maximal effort). The 0 on the scale (no effort) was anchored as the sensation associated with the participant being relaxed without contracting their muscle during 1 min. The 10 on the scale (maximal effort) was anchored as the sensation associated with the participant performing an MVC for 10 s. Participants were asked to consider the anchoring of ratings 0 and 10 when providing their RPE score during the experiment. Participants were also asked to not include any sensation of muscle pain or discomfort when rating their effort and to consider solely the sensation of “how hard it was to drive their leg” (Pageaux, 2016).

### Experiment 1: Torque Matching

Contraction type overview for the experiments is represented in Figure 2. Submaximal (20%, 40%, 60%, and 80% MVC) and maximal (100% MVC) voluntary contractions were randomly performed and matched at their given magnitude. Torque level was not disclosed to participants prior to beginning contractions. Torque and EMG were recorded during the trial. RPE was recorded immediately following the contraction by asking the participant the intensity of perceived effort experienced during the torque-plateau phase of their contraction. In order to minimize the possible confounding effects of muscle fatigue, the rTD (following active muscle shortening) contraction was performed first and the rTE (following active muscle lengthening) contraction was performed second. The active shortening muscle trial (rTD) consisted of a 2 s isometric pre-activation phase at −10° plantar flexion at the indicated torque level (% MVC) followed by an isokinetic (15° / s) shortening (10° dorsiflexion to 20° plantar flexion) over a 2 s period and ending with a 6 s steady state isometric phase at 20° plantar flexion. The active lengthening muscle trial (rTE) consisted of a 2 s isometric pre-activation phase at 50° plantar flexion at the indicated torque level followed by an isokinetic (15° / s) lengthening (50° to 20° plantar flexion) over a 2 s period and ending with a 6 s steady state isometric phase at 20° plantar flexion. A total of 10 contractions (five each for rTD and rTE states) were performed. Participants were given visual feedback of all torque traces on a computer monitor for all contractions. A 5 min rest period followed each contraction, as well as loosening of the inelastic straps about the ankle and metatarsals during periods of rest. Once all experimental contractions were completed, an MVC was performed and VA was reassessed to gauge the level of muscle fatigue.

**Fig. 2.**
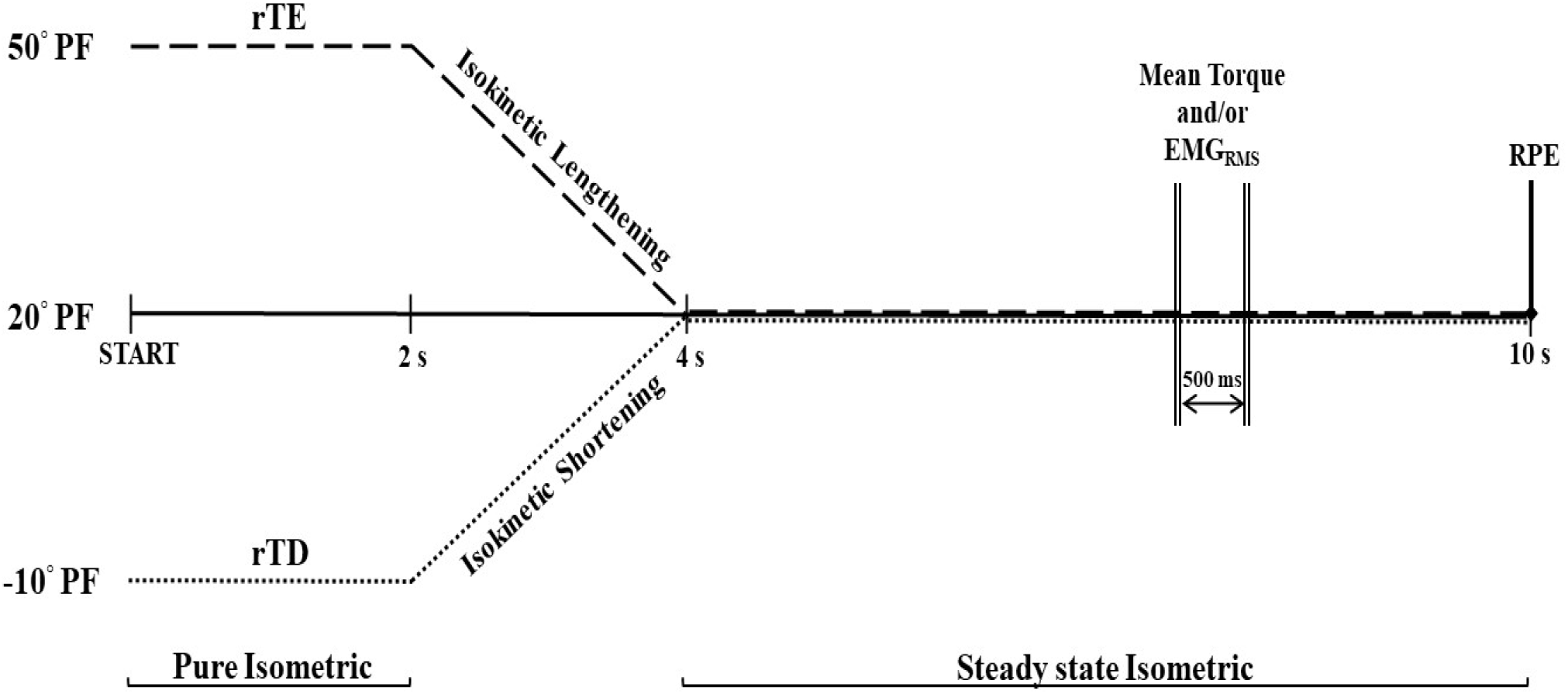
Schematic representation ofcontraction conditions during the experimental protocols. The rTD condition was first, where the contraction began at an ankle angle of 50° plantar flexion (PF) and the ankle was held at this angle for 2 s. Over the next 2 s, the ankle underwent isokinetic shortening at 15° / s to reach an ankle angle of 20° PF. The last 6 s of the contraction occurred at the 20° PF angle. The rTE trial followed the rTD trial, where the contraction began at an ankle angle of −10° PF and the ankle was held at this angle for 2 s. Over the next 2 s, the ankle underwent isokinetic lengthening at 15° / s to reach an ankle angle of 20° PF. The last 6 s of the contraction occurred at the 20° plantar flexion angle. EMG_RMS_ and/or torque were measured within a 500 ms window at the 7.5 – 8 s mark. RPE was taken immediately after the entire 10 s contraction for rTD and rTE trials.

### Experiment 2: Activation (iEMG) Matching

This experimental procedure used active shortening and lengthening contractions to induce a lower and higher torque output in the rTD and rTE states, respectively, while keeping EMG matched and recording participant RPE during the steady state isometric contraction. Integrated EMG (iEMG) was matched to a calculated ± 5% target window about the submaximal 20% and 80% maximal iEMG magnitude. Contraction intensity was randomized for each participant and was not disclosed to participants prior to the beginning of the contractions. The rTD state (active muscle shortening) contraction was performed first and the rTE state (active muscle lengthening) contraction was performed second. The rTD contraction consisted of a 2 s isometric pre-activation phase at −10° plantar flexion at the required iEMG level followed by an isokinetic (15° / s) shortening (10° dorsiflexion to 20° plantar flexion) over a 2 s period and ending with a 6 s steady state isometric phase at 20° plantar flexion. The rTE contraction consisted of a 2 s isometric pre-activation phase at 50° plantar flexion at the required iEMG level followed by an isokinetic (15° / s) lengthening (50° to 20° plantar flexion) over a 2 s period and ending with a 6 s steady state isometric phase at 20° plantar flexion. RPE was taken immediately following each contraction (rTD and rTE states). A total of four contractions were performed, one of each contraction type (rTD and rTE) for each trial at 20% and 80% iEMG. Participants were given visual feedback of the iEMG trace on a computer monitor for all contractions. A 5 min rest period followed each contraction, as well as loosening of the inelastic straps over the ankle and metatarsals during periods of rest.

### Experiment 3: Perception of Effort Matching

This experimental procedure was used to corroborate findings from experiments 1 and 2, to show no change in EMG amplitude during the rTD and rTE state whilst the intensity of perceived effort was held constant. Participants were asked to hold their perceived effort at a constant level during two trial conditions (RPE = 2, light intensity; RPE = 8, strong intensity) while both torque and EMG were recorded. The rTD state (active muscle shortening) contraction was performed first and the rTE state (active muscle lengthening) contraction was performed second. Participants performed the same contraction types in the same order for both light and strong intensities. The order of contraction intensities was randomized for both rTD and rTE contraction states. The rTD contraction consisted of a 2 s isometric pre-activation phase at 10° dorsiflexion, followed by an isokinetic (15° / s) shortening (−10° to 20° plantar flexion) over a 2 s period and ending with a 6 s steady state isometric phase at 20° plantar flexion. The rTE contraction consisted of a 2 s isometric pre-activation phase at 50° plantar flexion, followed by an isokinetic (15° / s) lengthening (50° to 20° plantar flexion) over a 2 s period and ending with a 6 s steady state isometric phase at 20° plantar flexion. Participants were given visual feedback of their torque trace on a computer monitor for all contractions. A 5 min rest period followed each contraction, as well as loosening of the inelastic straps on the ankle and metatarsals during periods of rest. Once all experimental contractions were completed, an MVC was performed and VA was reassessed to rule out any effects of muscle fatigue.

### Data Analysis

All participant data were analyzed with Labchart (Labchart, Pro Modules 2014, version 8) software. Torque (N·m), angular position (deg), and stimulus trigger signals were recorded and sampled at 1000 Hz using a 12-bit analog-to-digital converter (PowerLab System 16/35, ADInstruments, Bella Vista, AUS). Surface EMG data were sampled at 2000 Hz for both the soleus and tibialis anterior. Mean isometric torque and EMG_RMS_ amplitude were calculated in a 500 ms period at 7.5 – 8 s following the start of each contraction. In order to compare participant muscle activation between the rTD and rTE conditions, raw EMG was normalized (normalized EMG_RMS_) for each participant by dividing the EMG_RMS_ by their respective resting M_max_ (EMG_RMS_ / M_max_).

### Statistical Analysis

A paired samples t-test was performed on pre- and post-tests MVC peak torque, MVC peak EMG, and voluntary activation levels for all experiments. A student’s independent sample t-test was used to compare participant averages for each neuromuscular test between participants in experiment 1, and participants in experiments 2 and 3. For experiment 1, a 2 × 5 repeated measures ANOVA was used to assess the effects of history dependence (rTD and rTE states) and contraction intensity (20%, 40%, 60%, 80%, and 100% MVC) based on normalized EMG_RMS_ and RPE. For experiment 2, a 2 × 2 repeated measures ANOVA was used to assess the effects of history dependence (rTD and rTE states) and contraction intensity (20% and 80% iEMG_MVC_) on torque and RPE. For experiment 3, a 2 × 2 repeated measures ANOVA was used to assess the effects of history dependence (rTD and rTE states) and contraction intensity (light and strong effort) on torque and normalized EMG_RMS_ amplitude. In each experiment, the intensity was either controlled by torque (experiment 1), EMG (experiment 2), or the intensity of perceived effort (experiment 3). As our study aimed to compare the rTD and rTE states, all follow-up tests were performed solely in comparing these two history dependence states and the Bonferroni correction was applied. Significance was set at α = 0.05 (2-tailed) for all analysis. The effect sizes for the repeated measures ANOVAs were calculated as partial eta squared (η_p_^2^), and the paired-test comparison as Cohen’s d_z_. Descriptive data in the text and tables are reported as mean ± standard deviation (SD), and data reported in the figures are mean ± 95% confidence interval (CI). The paired samples t-test, independent student’s t-test, and fully repeated ANOVA were performed using jamovi (Version 1.2.27, Sydney, AUS). η_p_^2^ were obtained with jamovi software and Cohen’s d_z_ was with G*Power (Version 3.1.9.7, Düsseldorf, DEU).

## Results

Table 1 presents the between-experiments comparison in torque production capacity and neuromuscular function. There was no difference between participants (all p > 0.139). When comparing neuromuscular values pre- and post-experimental intervention, MVC peak torque (experiment 1: p = 0.634, d = 0.157; experiments 2 and 3: p = 0.780, d = 0.113), MVC peak EMG (experiment 1: p = 0.892, d = 0.019; experiments 2 and 3: p = 0.809, d = 0.045), or voluntary activation level (experiment 1: p = 0.327, d = 0.342; experiments 2 and 3: p = 0.213, d = 0.425) were not different. These results indicate that muscle fatigue did not influence our results.

### Experiment 1: Torque Matching

#### EMG

The main effect of contraction intensity revealed an increased normalized EMG_RMS_ amplitude (p < 0.001, η_p_^2^ = 0.745) when contraction intensity increased. The main effect of history dependence (p < 0.001, η_p_^2^ = 0.760) revealed that the normalized EMG_RMS_ amplitude was higher in the rTD state than the rTE state. There was an interaction between history dependence and torque (p = 0.006, η_p_^2^ = 0.155) where normalized EMG_RMS_ was lower in the rTD state than in the rTE state for matched torque intensities at 40% MVC (p < 0.001, d_z_ = 0.80), 60% MVC (p < 0.001, d_z_ = 0.79), 80% MVC (p < 0.001, d_z_ = 0.89), and 100% MVC (p < 0.001, d_z_ = 1.07). Results are presented in Figure 3A.

**Fig. 3.**
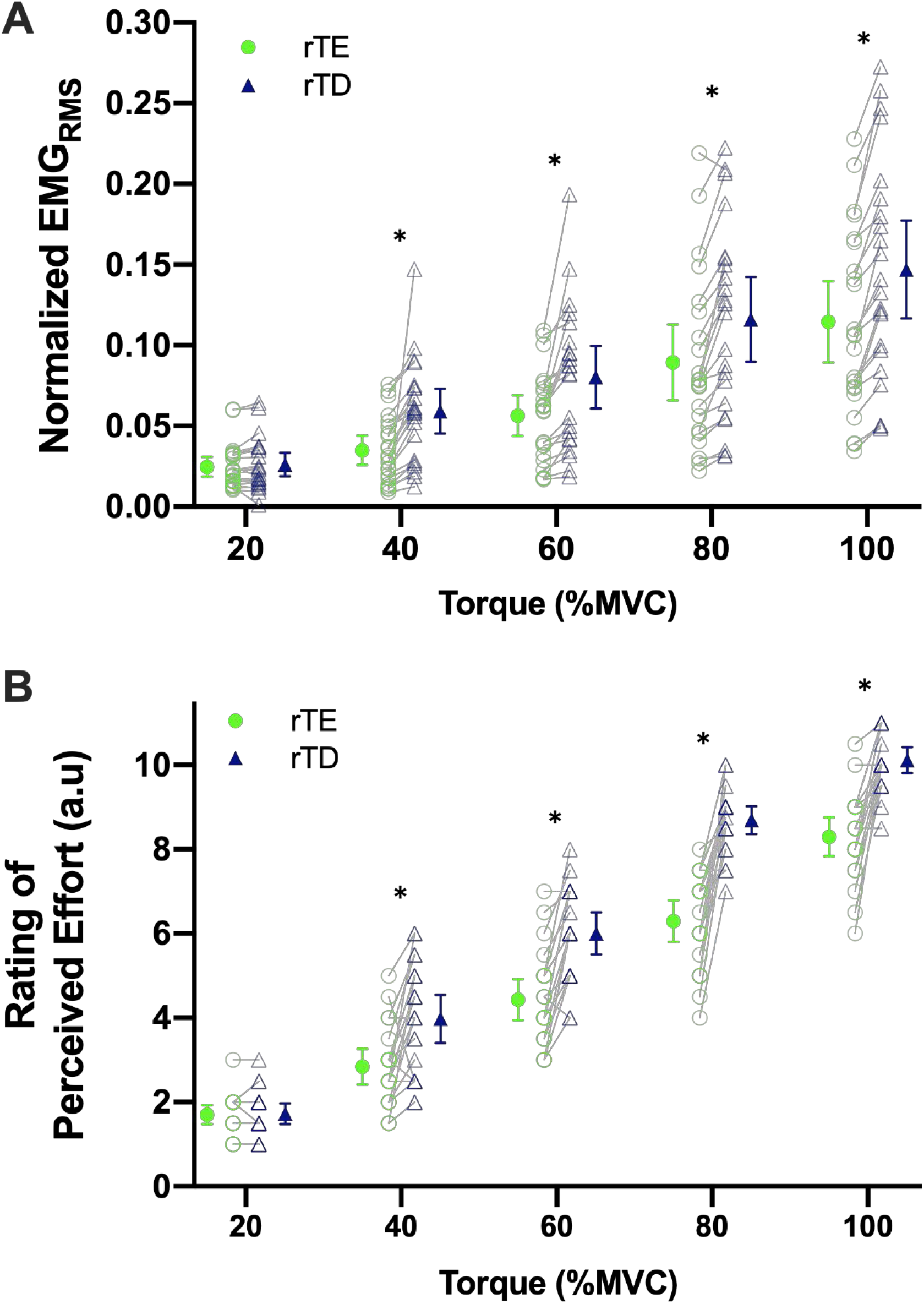
Experiment 1: Normalized EMG_RMS_ (panel A) and rating of perceived effort (panel B) when torque was matched. (A) There was an interaction between history dependence and torque contraction intensity (p = 0.006, η_p_^2^ = 0.155) for Normalized EMG_RMS_ between the rTD (triangle markers) and rTE (circle markers) states for torque matching intensities at: 40% MVC p < 0.001, 60% MVC p < 0.001, 80% MVC p < 0.001, and 100% MVC p < 0.001. There was no difference (p = 0.589) between the rTD and rTE normalized EMG_RMS_ at 20% MVC torque intensity. (B) There was an interaction between history dependence and torque contraction intensity (p < 0.001, η_p_^2^ = 0.246) for rating of perceived effort (RPE) between the rTD (blue) and rTE (green) states for torque matching intensities at: 40% MVC p < 0.001, 60% MVC p < 0.001, 80% MVC p < 0.001, and 100% MVC p < 0.001. There was no difference (p = 1) between the rTD and rTE RPE at 20% MVC torque intensity. Individual (N = 22) data are represented as empty markers and mean (±95%CI) as filled markers. Individual data with overlapping values can be identified with darker borders thickness. **\*** p < 0.001.

#### Perception of Effort

The main effect of contraction intensity revealed an increased RPE amplitude (p < 0.001, η_p_^2^ = 0.951) when contraction intensity increased. The main effect of history dependence (p < 0.001, η_p_^2^ = 0.831) revealed that the RPE was higher in the rTD state than in the rTE state. There was an interaction between history dependence and RPE (p < 0.001, η_p_^2^ = 0.246) where RPE was lower in the rTD state than in the rTE state for torque intensities at 40% MVC (p < 0.001, d_z_ = 1.52), 60% MVC (p < 0.001, d_z_ = 1.05), 80% MVC (p < 0.001, d_z_ = 1.60), and 100% MVC (p < 0.001, d_z_ = 1.22). Results are presented in Figure 3B.

### Experiment 2: iEMG Matching

#### Torque

The main effect of contraction intensity revealed an increased torque amplitude (p < 0.001, η_p_^2^ = 0.949) when contraction intensity increased. The main effect of history dependence (p < 0.001, η_p_^2^ = 0.962) revealed that torque was lower in the rTD state than in the rTE state. There was an interaction between history dependence and torque (p = 0.003, η_p_^2^ = 0.469) where torque was lower in the rTD state than the rTE state at 20% iEMG (p < 0.001, d_z_ = 1.28) and 80% iEMG (p < 0.001, d_z_ = 1.17). Results are presented in Figure 4A.

**Fig. 4.**
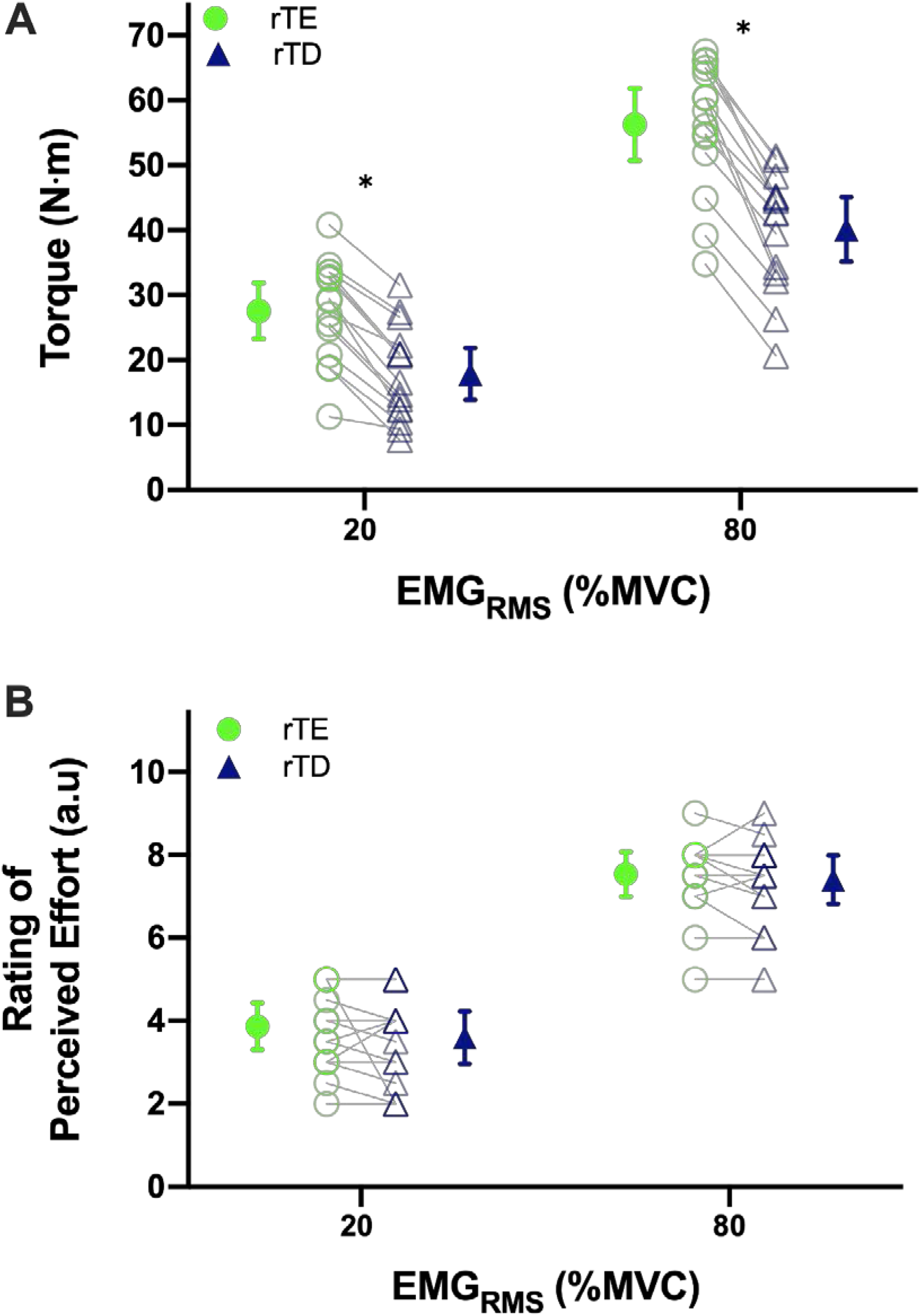
Experiment 2: Torque (panel A) and rating of perceived effort (panel B) when iEMG was matched. **(A)** There was an interaction between history dependence and EMG contraction intensity (p = 0.003, η_p_^2^ = 0.469) for torque between the rTD (triangle markers) and rTE states (circle markers) for all iEMG matching intensities (20% iEMG p < 0.001 and 80% iEMG p < 0.001). (B) There was no interaction between history dependence and EMG intensity (p = 0.541, η_p_^2^ = 0.027) for rating of perceived effort between the rTD and rTE states for all iEMG matching intensities. Individual (N = 15) data are represented as empty markers and mean (±95%CI) as filled markers. Individual data with overlapping values can be identified with darker borders thickness. * p < 0.001.

#### Perception of Effort

The main effect of contraction intensity revealed an increased RPE (p < 0.001, η_p_^2^ = 0.941) when contraction intensity increased. There was no main effect of history dependence (p = 0.228, η_p_^2^ = 0.102) revealing that RPE did not differ between the rTD state and the rTE state. The history dependence × RPE interaction did not reach significance (p = 0.541, η_p_^2^ = 0.027). Results are presented in Figure 4B.

### Experiment 3: Perception of Effort Matching

#### Torque

The main effect of contraction intensity revealed an increased torque (p < 0.001, η_p_^2^ = 0.859) when effort intensity increased. The main effect of history dependence (p < 0.001, η_p_^2^ = 0.740) revealed that torque was higher in the rTD state than in the rTE state. There was an interaction between history dependence and torque (p = 0.021, η_p_^2^ = 0.513) where torque was lower in the rTD state compared to the rTE state for both light (RPE = 2; p < 0.001, d_z_ = 0.88) and strong (RPE = 8; p < 0.001, d_z_ = 0.94) effort intensities. Results are presented in Figure 5A.

**Fig. 5.**
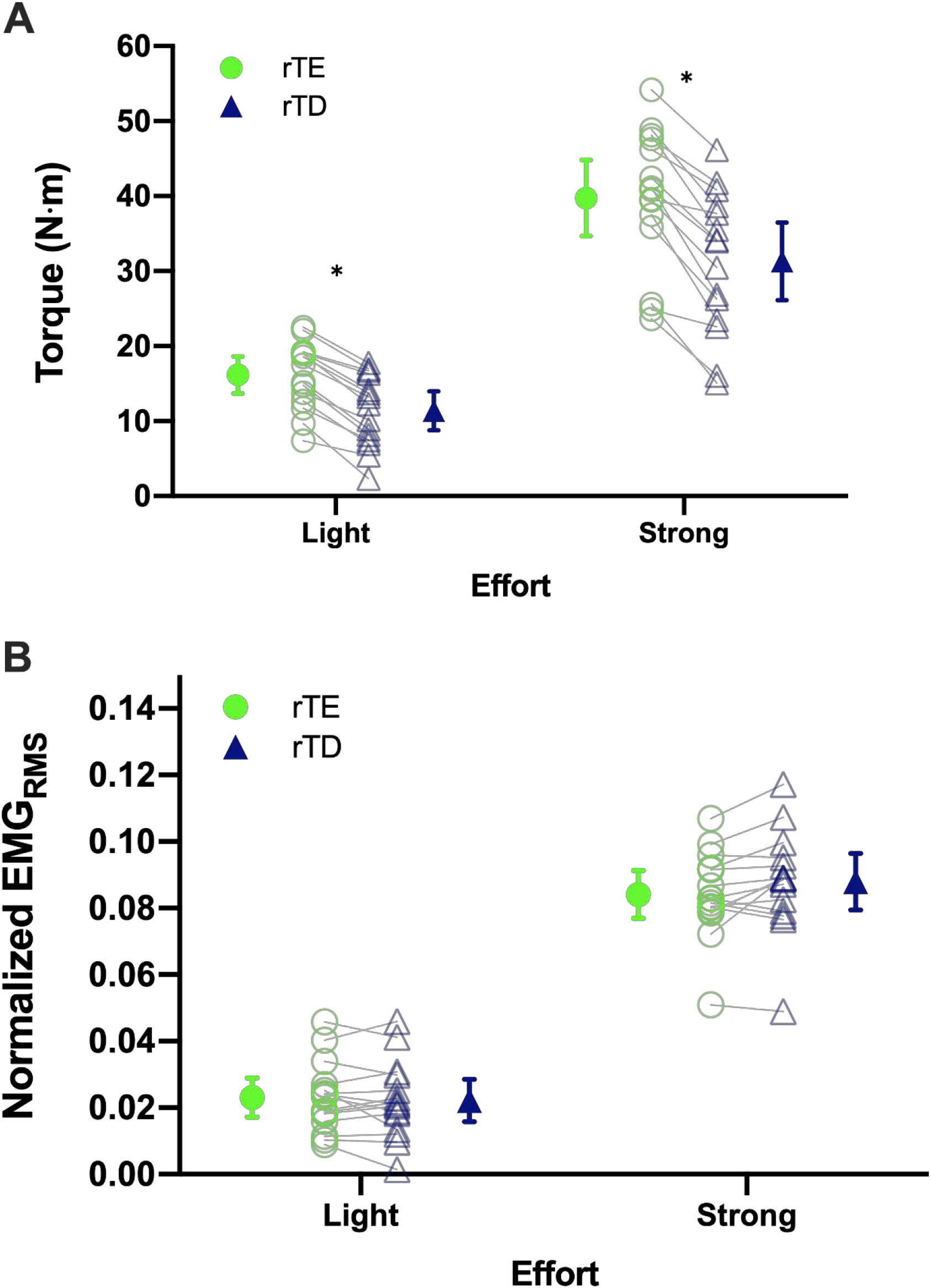
Experiment 3: Torque (panel A) and normalized EMG_RMS_ (panel B) when the intensity of perceived effort was matched. (A) There was an interaction between history dependence and the intensity of perceived effort (p = 0.021, η_p_^2^ = 0.513) for torque between the rTD and rTE state for both light (p < 0.001) and strong (p < 0.001) effort. (B) There was no interaction between history dependence and intensity of perceived effort (p = 0.366, η_p_^2^ = 0.059) for normalized EMG_RMS_ Individual (N = 15) data are represented as empty markers and mean (±95%CI) as filled markers. Individual data with overlapping values can be identified with darker borders thickness. **\*** p < 0.001.

#### EMG

The main effect of contraction intensity revealed an increased normalized EMG_RMS_ (p = 0.010, η_p_^2^ = 0.391) when the intensity of perceived effort increased. There was no main effect of history dependence (p = 0.354, η_p_^2^ = 0.062) revealing that the normalized EMG_RMS_ did not differ between the rTD state and the rTE state. The history dependence × EMG intensity interaction did not reach significance (p = 0.366, η _p_^2^ = 0.059). Results are presented in Figure 5B.

## Discussion

### History dependent states alter one’s perception of effort during the subsequent isometric contraction at matched torque

We investigated changes in the perception of effort evoked by rTE and rTD while matching torque and muscle activity (EMG). In experiment 1, torque matching conditions (40%, 60%, 80%, and 100% MVC) in the rTD state required greater neuromuscular activation whereas the rTE state showed a decrease in neuromuscular activation for all contraction intensities. The magnitude of activation increase or reduction was independent of torque production, which agrees with results from other studies that observed changes in EMG during history dependence of force (Chen et al., 2019; Jones et al., 2016; Paquin and Power, 2018; Seiberl et al., 2015). In line with our hypothesis, despite producing the same level of isometric steady state torque, one’s perception of effort is modulated in the isometric steady state following active muscle shortening or lengthening. When torque was matched, perception of effort was higher in the rTD state than in the rTE state for the 40%, 60%, 80%, and 100% MVC torque conditions, increasing in parallel with increased neuromuscular activation (EMG). We did not observe any difference in normalized EMG_RMS_ and perception of effort between the rTD and rTE states at a 20% MVC torque intensity. As neuromuscular activation assessed via EMG is traditionally used as a proxy of motor command sent to the working muscles (Carrier et al., 2011; Thoroughman and Shadmehr, 1999), the similar response in EMG and perception of effort at 20% MVC between the rTD and rTE state reinforces the hypothesis that perception of effort is dependent on the magnitude of motor command.

### Prior shortening and lengthening muscle actions do not alter perception of effort when neuromuscular activity is matched, despite producing distinct differences in torque

Results from experiment 2 indicate differing torque output in the isometric steady state following active muscle shortening and lengthening at the same iEMG level. The torque output was higher in the rTE state compared to the rTD state. Interestingly, despite a difference in torque output, perception of effort was closely matched with iEMG level. Indeed, no differences were observed in the RPE score between the rTD state and the rTE state when contraction intensity was matched with neuromuscular activation. As observed in experiment 1, the similar response in EMG and perception of effort across history-dependent states and contraction intensities reinforces the hypothesis that perception of effort is dependent on the motor command, and not the tension produced by the contracting muscle.

### Neuromuscular activation does not differ between steady state and purely isometric conditions when perceived effort is held constant, despite producing differing torque output

Results from experiment 3 indicate differing torque output in the isometric steady state following active muscle shortening and lengthening when held at the same level of perceived effort. When the intensity of perceived effort was matched in both light and strong contraction conditions, there was no difference in iEMG between the rTD and rTE states. However, the torque output was higher in the rTE state compared to the rTD state condition. As the torque output differed while neuromuscular activation was consistent between the two history dependence states, this experiment further reinforces the results of experiments 1 and 2, indicating that perception of effort is tightly coupled with the motor command.

### The physiological nature of history-dependent shortening and lengthening contractions influence perception of effort by modulating the magnitude of motor command, as reflected by the magnitude of neuromuscular activation recorded at the muscle

History dependent rTD and rTE muscle contractions produce distinct differences in EMG to match a given torque level (Herzog, 2004; Paquin and Power, 2018). During rTD, a larger EMG amplitude can be explained, at least in part, by altered cross-bridge mechanics after shortening. Marechal and Plaghki (1979) proposed that rTD could be associated with a stress-induced inhibition of cross-bridges in the newly formed actin-myosin overlap zone following shortening, straining, distorting, and inhibiting actin orientation of cross-bridge attachment sites. Direct inhibition of actin-myosin interaction results in fewer and less frequent cross-bridge formation (Herzog et al., 2000, 1998; Joumaa and Herzog, 2010; Pinnell et al., 2019). To overcome the mechanical limitations imposed by an actively shortened muscle, greater neuromuscular activation would be required to achieve equivalent isometric torque values, thus increasing the recruitment/firing rate of motor units. As such, the increased perception of effort during rTD can be, at least partially, explained by a greater motor command resulting in a higher neuromuscular activation as measured with EMG. In contrast, the rTE state elicits a decreased neuromuscular activation while torque remains constant (Gruber et al., 2009; Paquin and Power, 2018; Seiberl et al., 2012). Proposed mechanisms of rTE seem to stem from a greater contribution of passive force to overall total force production following active lengthening. It has been suggested that the Ca^2+^-dependent stiffening of the giant molecular spring, titin results in increased passive force following active lengthening (Joumaa and Herzog, 2014; Powers et al., 2016; Shalabi et al., 2017; Tahir et al., 2020). Due to the inherent mechanisms that contribute to greater torque during rTE at a matched EMG, less neuromuscular activation would be needed to maintain the same torque level. As such, the decreased perception of effort during rTE may be explained by a decrease in motor command. It seems that the magnitude of neuromuscular activation, and the perception of effort during the rTD and rTE states are tightly coupled even when absolute torque output is matched.

The origin for the perception of effort can be explained by the corollary discharge model, such that the efferent copy of generated motor commands are processed to generate the perception of effort (Christensen et al., 2007; de Morree et al., 2012; Marcora, 2009; Pageaux & Gaveau, 2016; Pageaux, 2016; Zénon et al., 2015). Movement related cortical potential, an electroencephalography marker of the motor command (de Morree et al., 2012; Jaquet et al. 2021) (REFs), and EMG amplitude have been shown to both increase during active muscle contractions. During muscle fatiguing tasks, both dynamic (de Morree et al., 2012) and isometric (Guo et al., 2017) muscle contractions have been shown to increase perceived effort and EMG, as well as larger movement related cortical potential amplitude, further substantiating that motor command influences perceived effort during active contraction. In addition, increased EMG is strongly related to increased perceived effort (Lampropoulou and Nowicky, 2012). During history dependent states, Seed et al., (2019) reported that perceived effort and EMG is distinctly higher in the rTD state compared to the rTE state when torque is matched at low, moderate, and high contraction intensities. Our study found a similar result for torque matching and extended these findings to activation and effort matching conditions. The difference in EMG between rTD and rTE states has been attributed to the motor command. Future investigation into changes of cortical responses between history dependent states (rTD and rTE) is warranted to substantiate a central origin to the perception of effort as mediated by the motor command, attributed to the phenomenon of the history dependence of force.

## Conclusions & Perspectives

In the present study, we attributed the ability of the history dependence of force (rTD or rTE state) to modulate perception of effort in the isometric steady state, to the magnitude of neuromuscular activation. Differences in perceived effort were observed between rTD and rTE states, with higher perceived effort reported in the rTD state due to the greater motor command needed to match the same absolute torque values. Accordingly, lower perceived effort was recorded in rTE state as a result of reduced motor command. When EMG was held constant between the rTD and rTE states, less force was produced in the rTD than in rTE state; however, as expected, perceived effort did not differ, thus supporting that perceived effort is determined by a central mechanism. This history dependence of force paradigm provides a useful model for the investigation of perceived effort during activation- or force-matched trials independent of time- and fatigue-dependent effects.

---

**Abbreviations.**
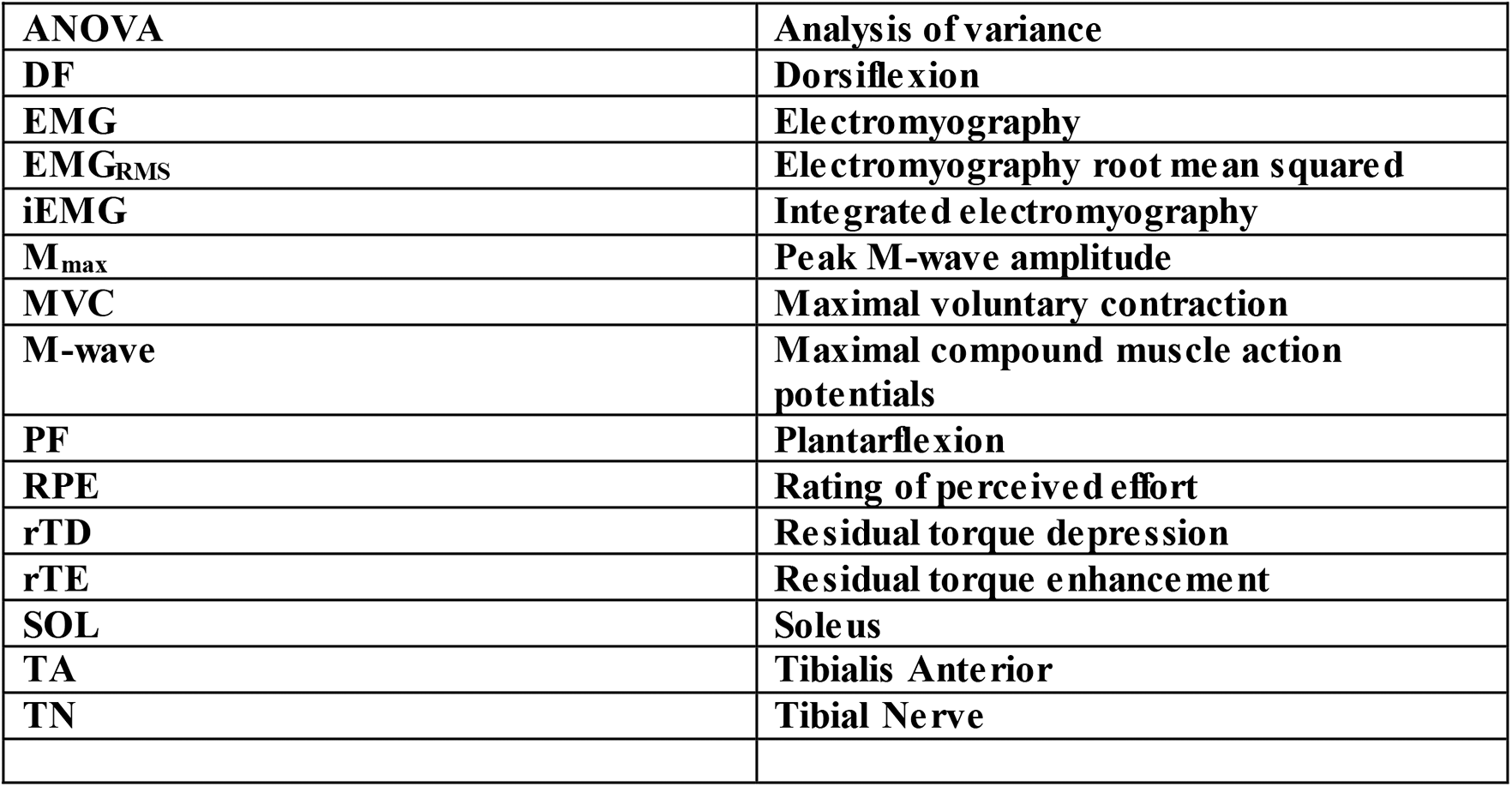

## Declarations

### Availability of data and material

Individual values of all supporting data are available upon request.

### Code availability

Not applicable

### Author contributions

BS, PJM, GAP designed the study. BP and GAP conducted the experiments. BK, BP, BS, GAP analyzed the data and formulated the results and figures. BK, BP, EFH, BS, PJM, GAP contributed to writing and editing the manuscript. All authors edited and approved the final submitted version of the manuscript.

### Ethics approval

Participants gave written informed consent prior to testing. All procedures were approved by the human Research Ethics Board of the University of Guelph (16-12-714).

### Consent to participate

Participants gave written informed consent prior to testing.

### Consent for publication

All authors edited and approved the final submitted version of the manuscript.

